# Detection of spreader nodes and ranking of interacting edges in Human-SARS-CoV protein interaction network

**DOI:** 10.1101/2020.04.12.038216

**Authors:** Sovan Saha, Piyali Chatterjee, Subhadip Basu, Mita Nasipuri

## Abstract

The entire world has recently witnessed the commencement of coronavirus disease 19 (COVID-19) pandemic. It is caused by a novel coronavirus (n-CoV) generally distinguished as Severe Acute Respiratory Syndrome Coronavirus 2 (SARS-CoV-2). It has exploited human vulnerabilities to coronavirus outbreak. SARS-CoV-2 promotes fatal chronic respiratory disease followed by multiple organ failure which ultimately puts an end to human life. No proven vaccine for n-CoV is available till date in spite of significant research efforts worldwide. International Committee on Taxonomy of Viruses (ICTV) has reached to a consensus that the virus SARS-CoV-2 is highly genetically similar to Severe Acute Respiratory Syndrome Coronavirus (SARS-CoV) outbreak of 2003. It has been reported that SARS-CoV has ∼89% genetic similarities with n-CoV. With this hypothesis, the current work focuses on the identification of spreader nodes in SARS-CoV protein interaction network. Various network characteristics like edge ratio, neighborhood density and node weight have been explored for defining a new feature spreadability index by virtue of which spreader nodes and edges are identified. The selected top spreader nodes having high spreadability index have been also validated by Susceptible-Infected-Susceptible (SIS) disease model. Initially, the proposed method is applied on a synthetic protein interaction network followed by SARS-CoV-human protein interaction network. Hence, key spreader nodes and edges (ranked edges) are unmasked in SARS-CoV proteins and its connected level 1 and level 2 human proteins. The new network attribute spreadability index along with generated SIS values of selected top spreader nodes when compared with the other network centrality based methodologies like Degree centrality (DC), Closeness centrality (CC), Local average centrality (LAC) and Betweeness centrality (BC) is found to perform relatively better than the *existing*-*state*-*of*-*art*.

## 1. Introduction

The pandemic COVID-19 registered its first case on 31 December 2019 [1]. It laid its foundation in the Chinese city of Wuhan (Hubei province) [2]. Soon, it made several countries all over the world [3] as its victim by community spreading which ultimately compelled World Health Organization (WHO) to declare a global health emergency on 30 January 2020 [4] for the massive outbreak of COVID-19. Owing to its expected fatality rate, which is about 4%, as projected by WHO [5], researchers from nations, all over the world, have joined their hands to work together to understand the spreading mechanisms of this virus SARS-CoV-2 [6]–[9] and to find out all possible ways to save human lives from the dark shadow of COVID-19.

Coronavirus belongs to the family Coronaviridae. This single stranded RNA virus not only affects humans but also mammals and birds too. Due to coronavirus, common fever/flu symptoms are noted in humans followed by acute respiratory infections. Nevertheless, corona viruses like Middle East Respiratory Syndrome (MERS) and Severe Acute Respiratory Syndrome (SARS) have the ability to create global pandemic due to their infectious nature. Both of these corona viruses are the member of genus Betacoronavirus under Coronaviridae. SARS created a major outbreak in 2003 originating from Southern China. 774 deaths were reported among 8098 globally registered cases resulting in an estimated fatality rate of 14%–15% [10]. While MERS commenced in Saudi Arabia creating an endemic in 2012. The world witnessed 858 deaths among 2494 registered positive cases. It generated high fatality rate of 34.4% in comparison to SARS [11].

SARS-CoV-2 is under the same Betacoronavirus genus as that of MERS and SARS coronavirus [12]. It comprises of several structural and non-structural proteins. The structural proteins includes the envelope (E) protein, membrane (M) protein, nucleocapsid (N) protein and the spike (S) protein. Though SARS-CoV-2 has been identified recently but there is an intense scarcity of data as well as necessary information needed to gain immunity against SARS-CoV-2. Studies has revealed the fact that SARS-CoV-2 is highly genetically similar to SARS-CoV based on several experimental genomic analyses [9], [12]–[14]. This is also the reason behind the naming of SARS-CoV-2 by International Committee on Taxonomy of Viruses (ICTV) [15]. Due to this genetic similarity, immunological study of SARS-CoV may lead to the discovery of SARS-CoV-2 potential drug development.

In the proposed methodology, Protein-protein interaction network (PPIN) has been used as the central component in identification of spreader nodes in SARS-CoV. PPIN has been found to be very effective module for protein function determination [16]–[18] as well as in the identification of central/essential or key spreader nodes in the network [19]–[23]. Compactness of the network and its transmission capability is estimated using centrality analysis. Anthonisse *et al.* [23] proposed a new centrality measure named as Betweenness Centrality (BC). BC is actually defined as the measure of impact of a particular node over the transmission between every pair of nodes with a consideration that this transmission is always executed over the shortest path between them. It is defined as:

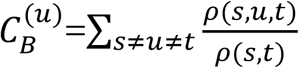

where *ρ*(*s, t*) is the total number of shortest paths from node *s* to node *t* and *ρ*(*s, u, t*) is the number of those paths that pass through *u*. Another centrality measure, called as closeness centrality (CC), was defined by Sabidussi *et al.* [24]. It is actually a procedure for detecting nodes having efficient transmission capability within a network. Nodes which have high closeness centrality value are considered to have the shortest distance to all available nodes in the network. It can be mathematically expressed as:

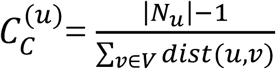

where |*N*_*u*_| denotes the number of neighbors of node u and *dist*(*u, v*) is the distance of the shortest path from node *u* to node *v*. Two other important centrality measures: Degree centrality (DC) [19] and Local average centrality (LAC) [21] are also found to be very effective in this area of research. DC is considered to be the most simplest among the available centrality measures which only count the number of neighbors of a node. Nodes having high degree is said to be highly connected module of the network. It is defined as:

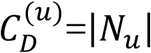

where |*N*_*u*_| hold the same meaning as state above. LAC is defined to be the local metric to compute the essentiality of node with respect to transmission ability by considering its modular nature, the mathematical model of which is highlighted as:

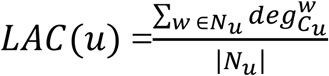

where *C*_*u*_ is the subgraph induced by *N*_*u*_ and 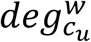 is the total number of nodes which are directly connected in *C*_*u*_.

Due to high morbidity and mortality of SARS-CoV2, it has been felt that there is a pressing need to properly understand the way of disease transmission from SARS-CoV-2 PPIN to human PPIN [16]–[18]. But since SARS-CoV-2 PPIN is still not available, SARS-CoV PPIN is considered for this research study due to its high genetic similarity with SARS-CoV-2. In the proposed methodology, at first SARS-CoV-Human PPIN (up to level 2) is formed from the collected datasets [25]–[27]. Once it is formed, spreader nodes are identified in each of SARS-CoV proteins, its level 1 and level 2 of human network by the application of a new network attribute i.e. spreadability index which is a combination of three terminologies: 1) edge ratio [28] 2) neighborhood density [28] and 3) node weight [29]. The detected spreader nodes are also validated by the existing SIS epidemic disease model [30]. Then the edges connecting two spreader nodes are ranked based on the average of spreadability index of spreader nodes themselves to access to the spreading ability of the corresponding edge. The ranked edges thus highlight the path of entire disease propagation from SARS-CoV to human level 1 and then from human level 1 to human level 2 proteins.

## 2. Methodology

Spreader nodes and edges both play a crucial role in transmission of infection from one part of the network to another. Generally in a disease specific PPIN models, at least two entities are involved: one is pathogen/Bait and the other is host/Prey [31]. In this research work, SARS-CoV takes the role of the former while human later. SARS-CoV first transmits infection to its corresponding interaction human proteins which in turn affect its next level of proteins. So, the transmission occurs through connected nodes and edges. Not all the nodes or proteins in the PPIN transmit infection. So, proper identification of nodes transmitting infection (spreader nodes) is required. It is simultaneously also true that the transmission is not possible without the edges connecting two spreader nodes. Thus these connecting edges are called spreader edges. The proposed methodology involves a proper study and assessment of various existing established PPIN features followed by the identification of spreader nodes, which has been also verified by SIS model.

### 2.1 Basic Network Characteristics

#### 2.1.1 Ego Network

Ego network of node *i* (*S*_*i*_) [28] is defined as the grouping of node *i* itself along with its corresponding level 1 neighbors and interconnections. N (*S*_*i*_) [28] consists of the set of nodes which belong to the ego network, *S*_*i*_ i.e. {*i*} ∪ *Γ*(*i*).

#### 2.1.2 Edge ratio

Edge ratio of node *i* [28] is defined by the following equation:

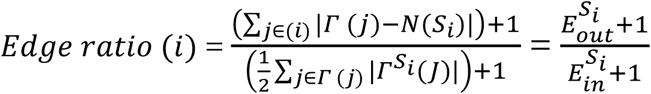

where 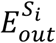 is the total number of outgoing edges from the ego network *S*_*i*_ and 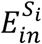 is the total number of interactions among node *i*’s neighbors, respectively. Γ (*i*) denotes the level 1 neighbors of node *i. S*_*i*_ is considered to be Ego network. 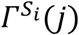 denotes node *j*’s neighbors which belongs *S*_*i*_. In the edge ratio, 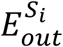 is positively related to the non-peripheral location of node *i*. Large number of interactions resulting from the ego network actually denotes that the node has high level of interconnectivity between its neighbors. On the other hand, 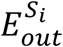 is negatively related to the inter-module location of node *i*. It represents the fact that the interconnectivity between neighbors is usually connected to the number of structural holes available around the node. When neighbor’s interconnectivity is low, the root or the central node *i* gain more control of the flow of transmission among the neighbors.

#### 2.1.3 Jaccard Dissimilarity

Jaccard dissimilarity [32] of node *i* and *j* (*dissimilarity* (*i, j*)) is defined as:

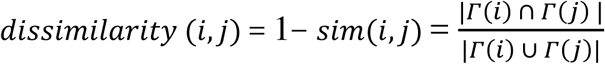

where |Γ(*i*) ∩ Γ(*j*) | refers to the number of common neighbors of neighbors of *i* and *j*. |Γ(*i*) ∪ Γ(*j*)| is the total number of neighbors of *i* and *j*. Similarity between two nodes is determined by Jaccard similarity based on their common neighbors. The similarity degree between *i* and *j* is considered to be more when they have more number of common neighbors. Whereas, when dissimilarity between the neighbors of a node is high, it guarantees that the only common node among the neighbors is the central node, which is termed as structural hole situation [28].

#### 2.1.4 Neighborhood Density

The neighborhood diversity [28] is defined as:

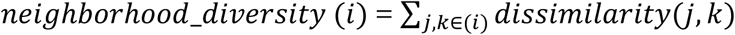

When a node’s neighborhood density reaches its greatest value, it reveals the fact that the neighbors have no other closer path. Hence, the neighbors should transmit or communicate through this node.

#### 2.1.5 Node weight

Node weight *w*_*v*_ of node *v* ∈ *V* in PPIN [29] is interpreted as the average degree of all nodes in *G*_*V*′_, a sub-graph of a graph *G*_*V*_. It is considered as another measure to determine the strength of connectivity of a node in a network. Mathematically, it is represented by

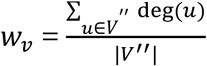

where, *V*” is the set of nodes in G_*V*′_. | *V*″| is the number of nodes in G_*V*′_. And *deg*(*u*) is the degree of a node *u* ∈ *V*”.

### 2.2 Spreadability Index and Spreader nodes

Spreadability index of node *i* is defined as the ability of node *i* to transmit an infection from one node to another. Mathematically it can be defined as:

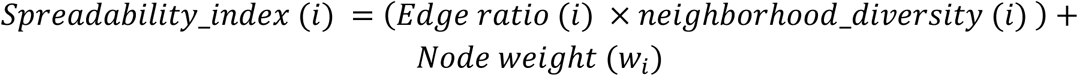

Nodes having high spreadability index are termed as spreader nodes i.e. these nodes have the capability to transmit the infection to its maximum number of interconnected neighbors in a much short amount of time in comparison to the other nodes in PPIN.

Figure 1 represents a sample PPIN where each protein is denoted as nodes while its interaction by edges. In this network, spreadability index is computed by using basic network characteristics as stated earlier which has been highlighted and compared to DC, BC, CC and LAC in Table 1 to Table 5.

**Table 1.** Computation of spreadability index of synthetic figure 1 along with computation of infection rate of selected top 10 spreader nodes by SIS model.

| Rank | Proteins | $E_{out}^{S_i}$ | $E_{in}^{S_i}$ | Edge Ratio | Neighborhood Density | Node Weight | Spreadability Index | Sum of SIS infection rate of top 10 nodes |
| --- | --- | --- | --- | --- | --- | --- | --- | --- |
| 1 | 1 | 13 | 0 | 14.0 | 5.19 | 3.40 | 76.15 | 2.46 |
| 2 | 24 | 4 | 2 | 1.66 | 12.5 | 1.87 | 22.70 |  |
| 3 | 4 | 6 | 1 | 3.50 | 3.63 | 2.40 | 15.11 |  |
| 4 | 5 | 8 | 3 | 2.25 | 4.39 | 3.60 | 13.48 |  |
| 5 | 19 | 6 | 2 | 2.33 | 3.8 | 2.80 | 11.66 |  |
| 6 | 23 | 5 | 4 | 1.20 | 6.58 | 3.00 | 10.89 |  |
| 7 | 17 | 4 | 1 | 2.50 | 3.33 | 2.00 | 10.33 |  |
| 8 | 6 | 7 | 0 | 8.00 | 0.87 | 3.00 | 10.00 |  |
| 9 | 2 | 4 | 4 | 1.00 | 6.84 | 2.83 | 9.68 |  |
| 10 | 22 | 6 | 4 | 1.40 | 3.88 | 3.60 | 9.03 |  |
| 11 | 25 | 7 | 0 | 8.00 | 0.71 | 3.00 | 8.71 | -- |
| 12 | 27 | 7 | 0 | 8.00 | 0.71 | 3.00 | 8.71 |  |
| 13 | 28 | 7 | 0 | 8.00 | 0.71 | 3.00 | 8.71 |  |
| 14 | 30 | 7 | 0 | 8.00 | 0.71 | 3.00 | 8.71 |  |
| 15 | 18 | 6 | 0 | 7.00 | 0.85 | 2.66 | 8.66 |  |
| 16 | 20 | 7 | 2 | 2.66 | 1.78 | 3.50 | 8.26 |  |
| 17 | 7 | 4 | 3 | 1.25 | 4.15 | 2.80 | 7.98 |  |
| 18 | 21 | 3 | 6 | 0.57 | 6.66 | 3.33 | 7.13 |  |
| 19 | 3 | 3 | 3 | 1.00 | 4.00 | 2.60 | 6.60 |  |
| 20 | 16 | 4 | 2 | 1.66 | 2.06 | 2.75 | 6.19 |  |
| 21 | 15 | 4 | 2 | 1.66 | 2.06 | 2.75 | 6.19 |  |
| 22 | 31 | 6 | 1 | 3.50 | 0.75 | 3.33 | 5.95 |  |
| 23 | 33 | 6 | 1 | 3.50 | 0.75 | 3.33 | 5.95 |  |
| 24 | 32 | 4 | 2 | 1.66 | 1.75 | 2.75 | 5.66 |  |
| 25 | 8 | 4 | 3 | 1.25 | 1.88 | 3.25 | 5.60 |  |
| 26 | 14 | 6 | 0 | 7.00 | 0.40 | 2.66 | 5.46 |  |
| 27 | 9 | 2 | 4 | 0.60 | 3.64 | 2.80 | 4.98 |  |
| 28 | 10 | 5 | 1 | 3.00 | 0.50 | 3.00 | 4.50 |  |
| 29 | 13 | 1 | 3 | 0.50 | 1.70 | 2.50 | 3.35 |  |
| 30 | 11 | 1 | 3 | 0.50 | 1.70 | 2.50 | 3.35 |  |
| 31 | 12 | 1 | 3 | 0.50 | 1.70 | 2.50 | 3.35 |  |
| 32 | 29 | 2 | 0 | 3.00 | 0.00 | 1.33 | 1.33 |  |
| 33 | 26 | 2 | 0 | 3.00 | 0.00 | 1.33 | 1.33 |  |

**Figure 1.**
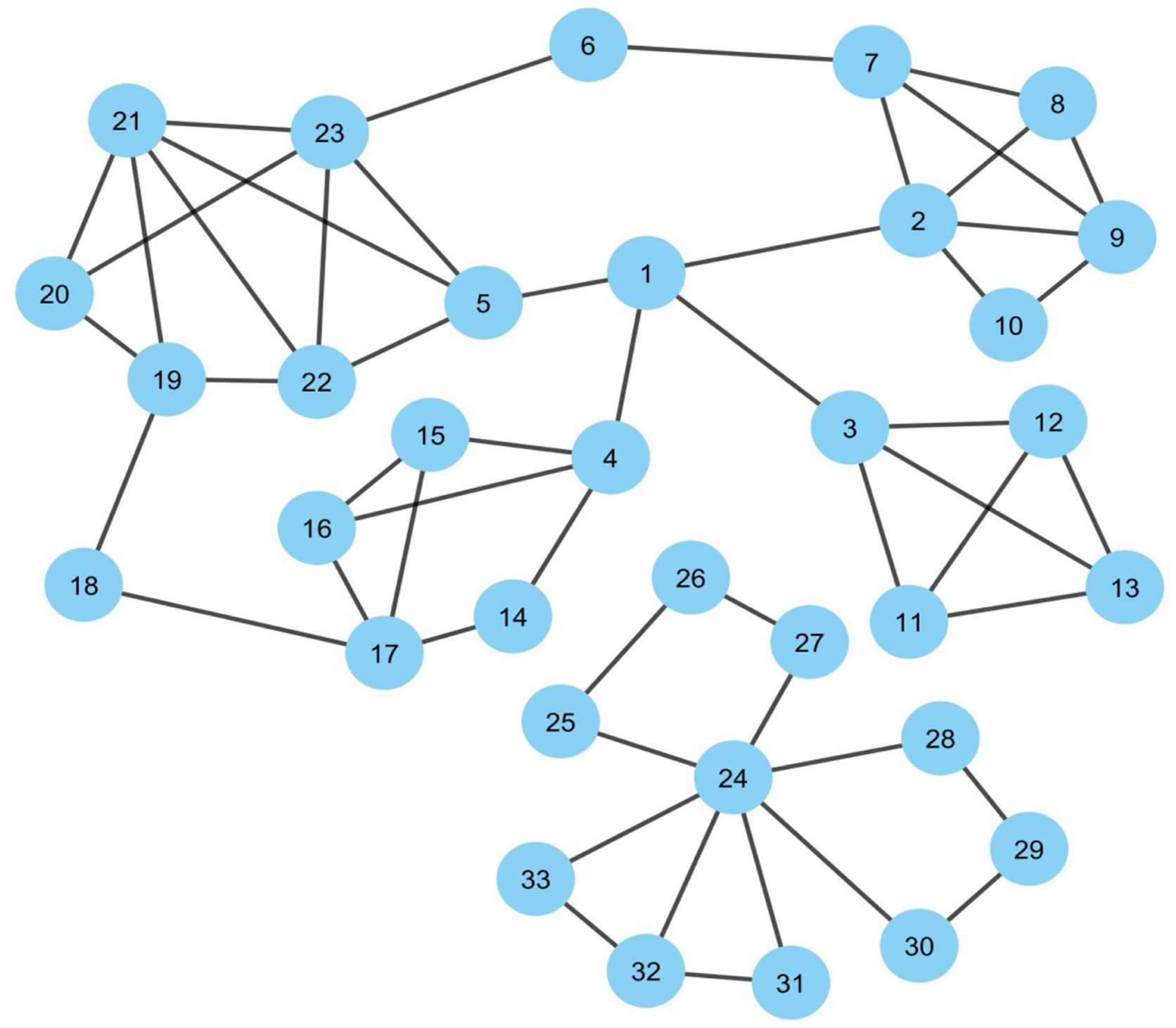
Synthetic PPIN1 consisting of 33 nodes and 53 edges where each node represents a protein and its interaction is represented by edges.

In Figure 1, it can be observed that nodes 1, 24 are clearly the most essential spreaders. Node 1 connects the four densely connected modules of the network which turns this node to stand in the first position having highest spreadability index by our proposed methodology. This node has been correctly ranked by all the methods except LAC and DC. Node 24 though has moderate edge ratio and node weight, but is one of the most densely connected modules itself in spite of getting isolated from the main network module of node 1. Moreover, node 24 has the highest neighborhood density. It establishes the fact that the only path of transmission for nodes 26, 27, 25, 28, 29, 30, 31, 32 and 33 is node 24. Thus if node 24 gets affected, then all the connected nodes with it will be immediately affected due to the lack of the connectivity of the neighbors with other central nodes. So, node 24 holds the second position with respect to spreadability index in our proposed methodology. Node 24 is not properly identified as the second most influential spreader nodes by the other methods. Further assessment of the remaining nodes highlights the fact that the performance of the new attribute spreadability index in our proposed methodology is relatively better in comparison to the others.

### 2.3 Validation of spreader nodes by SIS model

The SIS Epidemic Model [30] is used in this proposed methodology as a way of modeling SARS-CoV epidemic by classifying the proteins in SARS-CoV-human PPIN based on their disease/infection status. SIS actually refers to **S**usceptible, **I**nfected and **S**usceptible states which are generally considered as the three probable states of a protein in a PPIN. 1) **S** - The susceptible proteins who are not yet infected but are at risk for getting infected. In general every protein in PPIN is initially in susceptible state. 2) **I** - Infected proteins who are infected and are capable of transmitting the disease to other proteins. 3) **S** - proteins who have recovered and again become susceptible. Infection rate of the network, recovery rate (general assumption is any one out of the infected proteins gets recovered in one day) and total number of proteins are usually provided as input to SIS. If a protein gets infected and it has many neighbors, then any neighbor can get infected or may not be. So the final result is generated after 50 iterations for each infected protein. The total number of susceptible proteins after 50 iterations in the neighborhood of each infected protein divided by the total number of proteins in the network gives the infection capability of the infected protein. Thus the spreader nodes identified by spreadability index are validated by the infection rate as generated by SIS for them. It can be observed from Table 1 to Table 5 that the proposed methodology has the highest SIS infection rate of 2.46 (see Table 1) in comparison to others for their corresponding top 10 spreader nodes in the synthetic network as shown in Figure 1.

### 2.4 Ranking of Spreader edges

To show the ranking of interacting spreader edges, another synthetic PPIN 2 has been considered as network 1 while the previous synthetic PPIN 1 in section 2.2 (Figure 1) has been represented as network 2 in Figure 2. Node D, E and F are the selected top spreader nodes in network 1 by spreadability index in the same way as that of synthetic PPIN 1 in Figure 1 while to avoid the complexity in the diagram, top 5 nodes in network 2 (see Table 1) are selected as spreader nodes in network 2. Red colored edges are the interconnectivities between the nodes of network 1 while black colored edges show the interconnectivity between nodes of network 2. Green colored spreader edges (i.e. edges connected with spreader nodes) show the interconnectivity between network 1 and 2. Ranking of a spreader edge measures the effectiveness of transmission ability of a spreader edge i.e. how many interactive nodes gets infected through that edge. Thus all the spreading edges are ranked based on the average of the spreadability index of its connected spreader nodes. The ranked spreader edges in Figure 2 are highlighted in Table 6.

**Figure 2.**
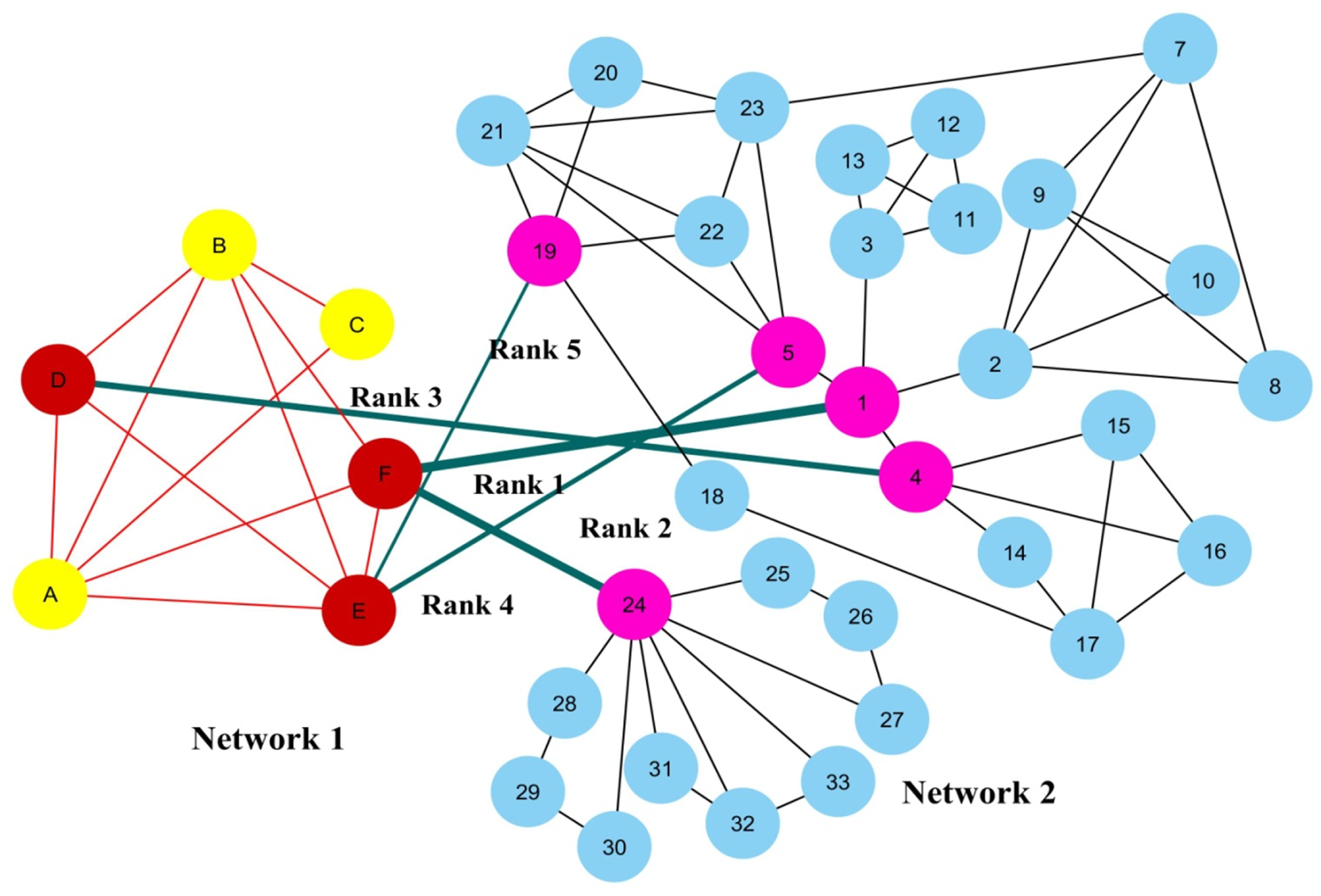
Ranking of spreader edges between two interconnected PPIN based on spreadability index. Thickness of the edges vary with the order of ranking.

## 3. Experimental Result

The proposed methodology leads to the identification of spreader nodes and edges through a network characteristic, called spreader index which has been also checked and validated by SIS model. Initially the whole working module is implemented on synthetic networks as shown in Figure 1 and Figure 2 in section 2 and then on SARS-CoV-human dataset. To visualize the infection transmission from SARS-CoV to human, PPIN of the both needs to be formed for which three datasets have been curated. They are: 1) SARS-CoV PPIN [25] 2) SARS-CoV-Human PPIN [25] 3) Human PPIN [26], [27]. After the removal of self-loops and data redundancy, the final SARS-CoV PPIN consists of 17 interactions among SARS-CoV proteins along with the involvement of 7 unique proteins after the removal of self loops and data redundancy. Only the densely interconnected SARS-CoV proteins having direct connections (level 1) with human proteins are considered rather than the isolated ones. SARS-CoV-Human PPIN includes 118 interactions between SARS-CoV and human. It is used to fetch the level 1 interaction of human proteins for the corresponding SARS-CoV proteins in SARS-CoV PPIN. Human PPIN consists of 314384 interactions. It is utilized for getting the indirect interactions (level 2) of level 1 human proteins formed earlier. The application of the proposed methodology in SARS-CoV-human PPIN is highlighted in Figure 3.

**Figure 3.**
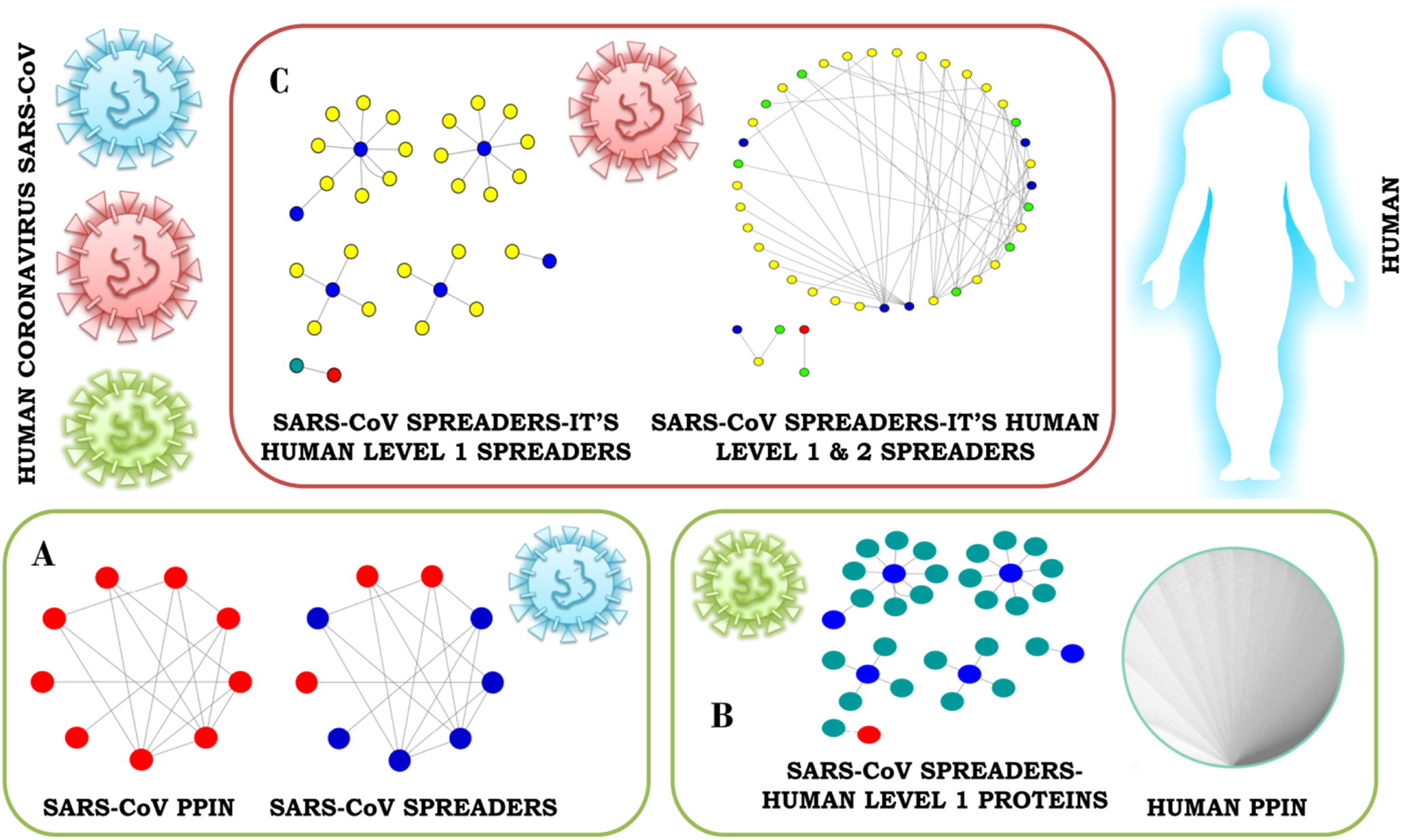
Mechanism of transmission of infection through spreader nodes. **A**. PPIN of SARS-CoV (red). Selection of spreader nodes in SARA-CoV PPIN (blue). **B.** Interaction of SARS-CoV spreaders with its level 1 corresponding proteins in human PPIN (green). **C.** Selection of spreaders in level 1 (yellow) and level 2 human proteins (green). Rest proteins in human PPIN are ignored to prevent overlap in the diagram.

In Figure 3A, at first SARS-CoV PPIN is displayed in which each protein is marked in red. Thereafter spreader nodes in SARS-CoV PPIN are identified by spreadability index which are denoted as blue nodes among red. Once the spreader nodes are active (in Figure 3B), it transmits the infection to its corresponding direct partners i.e. human level 1proteins (marked in green). In Figure 3C, spreader nodes are identified in SARS-CoV level 1 human proteins (marked in blue) and the disease will be transmitted further. The same will continue till SARS-CoV level 2 human proteins in which green nodes are the spreaders and thus the infection will penetrate further in human PPIN resulting in a significant fall in human immunity level followed by severe acute respiratory syndrome.

In Figure 4, the PPIN of SARS-CoV network has been highlighted. There are mainly 9 proteins which include E, M, ORF3A, ORF7A, S, N, ORF8A, ORF8AB, and ORF8B. The computed spreadability index of each of this protein and its corresponding validation by SIS model is highlighted in Table 7. It is also compared with other central/ influential spreader node detection methodologies like DC, CC, LAC and BC which are shown in Table 8, Table 9, Table 10 and Table 11 respectively. Similarly spreader nodes are also identified in SARS-CoV’s level 1 neighbors and level 2 neighbors (see Figure 5 and Figure 6).

**Figure 4.**
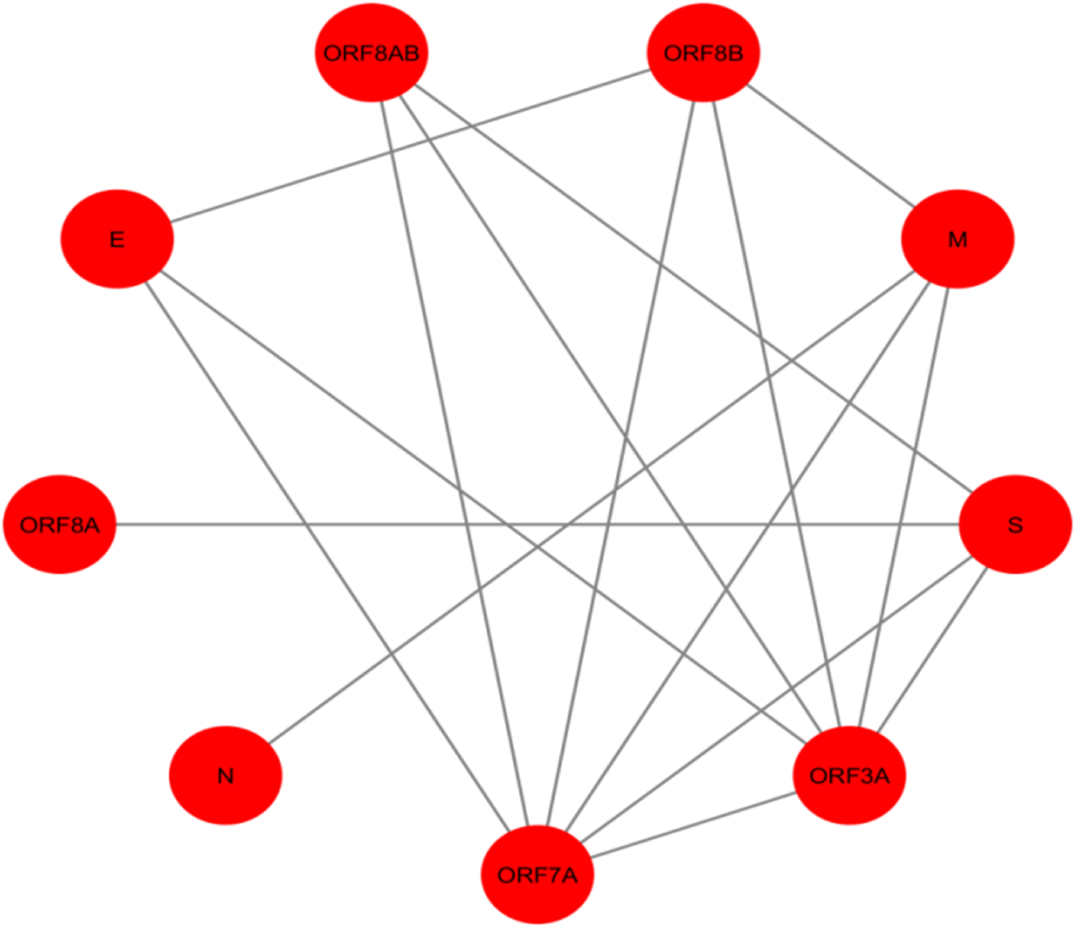
SARS-CoV PPIN consisting of 9 proteins which each protein is represented by a node and its corresponding interaction is highlighted by edges.

**Figure 5.**
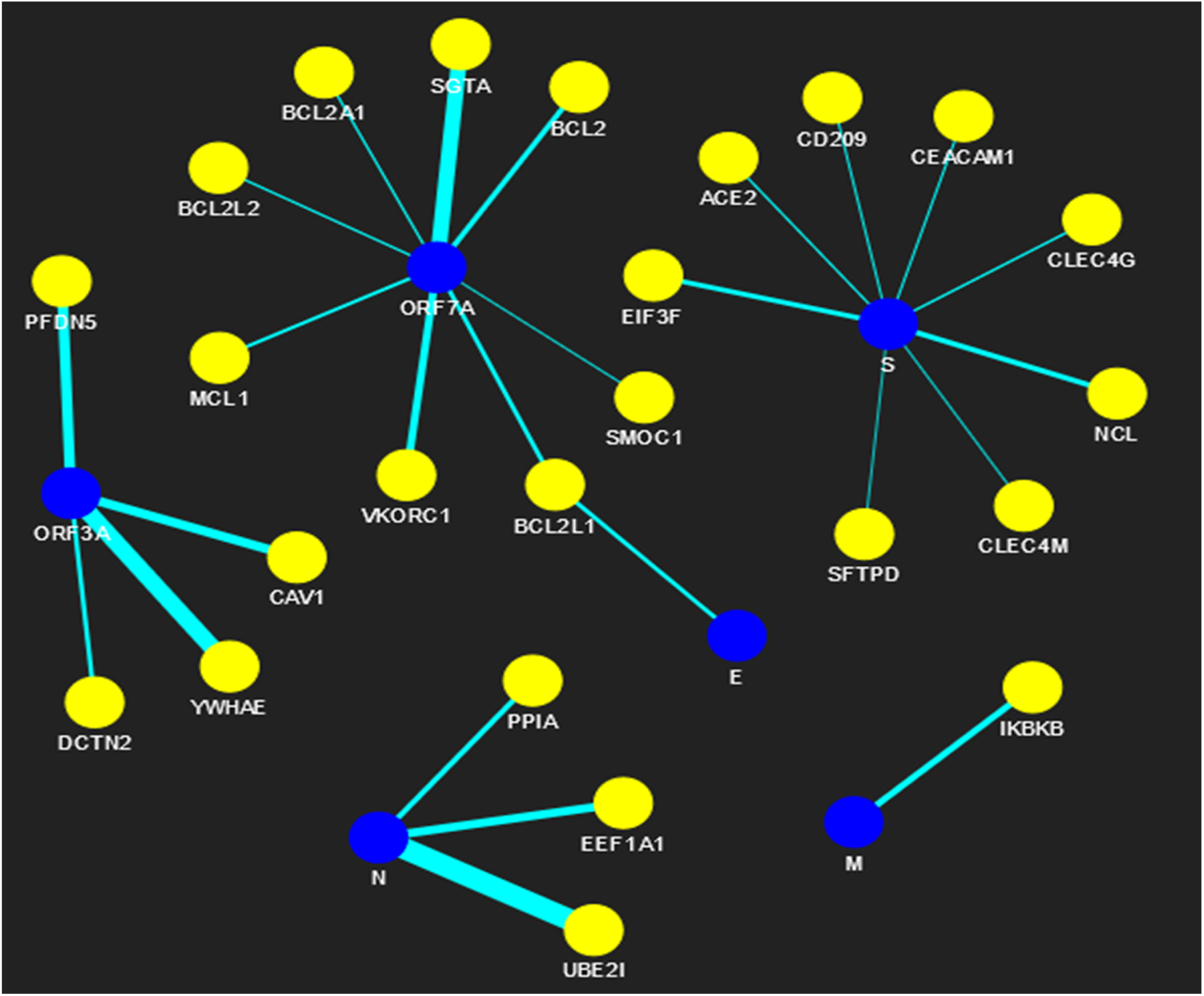
SARS-CoV-human PPIN (blue node represents SARS-CoV proteins while yellow node represents SARS-CoV s level 1 human spreaders)

**Figure 6.**
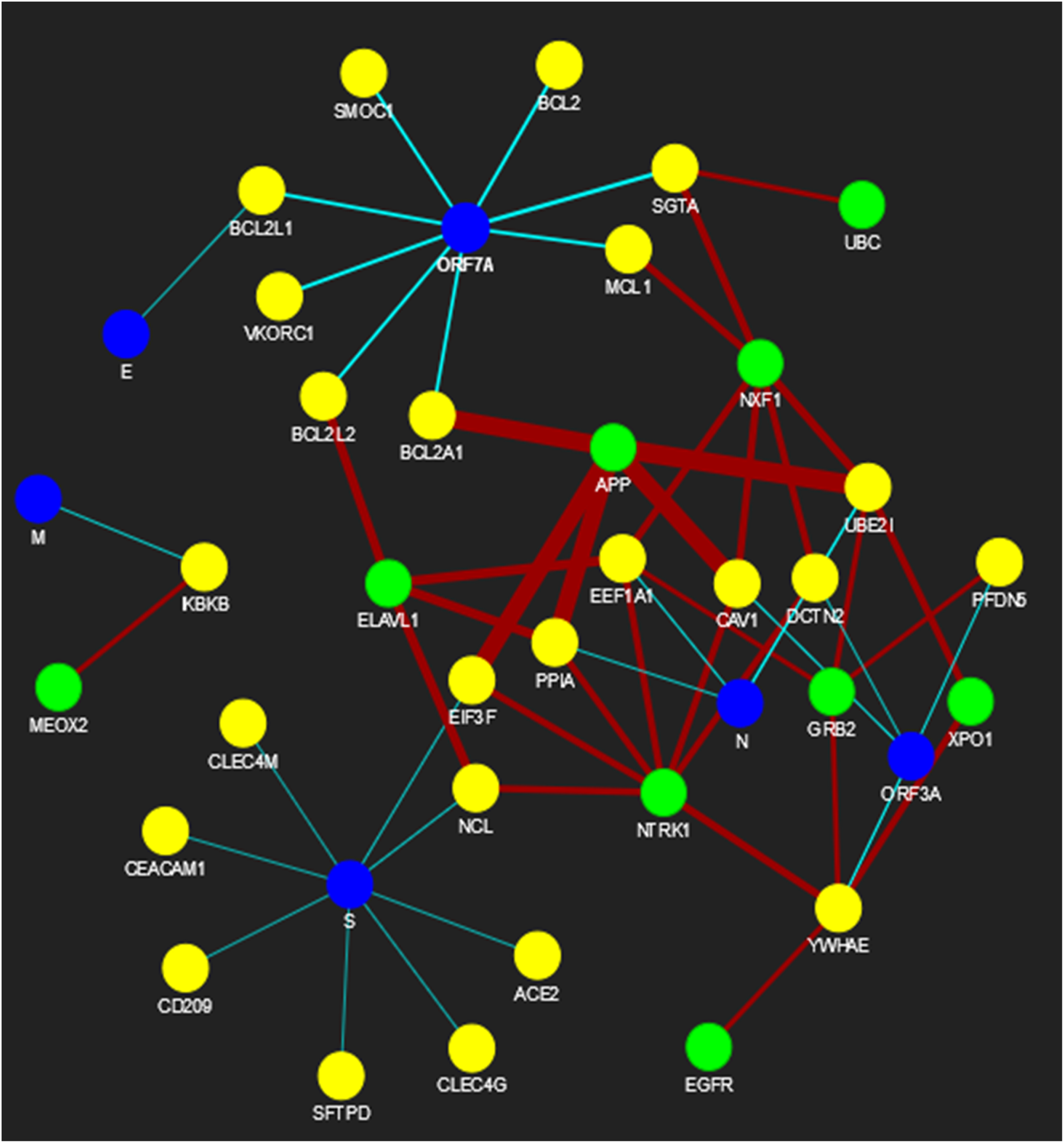
SARS-CoV-human PPIN (blue node represents SARS-CoV proteins while yellow and green nodes represent SARS-CoV s level 1 and level 2 human spreaders respectively)

Spreadability index definitely plays an important role in this proposed methodology. In fact, spreader nodes are successfully identified by the virtue of this scoring technique which covers all the aspects of transmitting infection from one node to another in a PPIN. It should be mentioned here that while identifying spreader nodes in SARS-CoV level 2 human proteins, it has been noted that the number of nodes are getting increased largely with the increment of successive levels. So, high, medium and low threshold [33] have been applied and the entire disease transmission through spreadability index is computationally assessed at each threshold.

The network statistics of spreader nodes at each level of threshold is shown in Table 12. It can be observed that threshold application is only implemented at SARS-CoV level 2 human proteins not on others. This is because of the availability of very less number of nodes and edges. Only nodes and edges having extremely low spreadability index have been discarded at the first level.

Beside identification of spreader nodes, spreader edges are also identified. The ranked edges between SARS-CoV spreaders and its level 1 human spreaders is highlighted in Table 13 while the ranked edges between SARS-CoV s level 1 and level 2 human spreaders at high, medium and low threshold are highlighted in A-Table 1, A-Table 2 and A-Table 3 in the appendix respectively.

**Table 2.** Computation of CC of synthetic figure 1 along with computation of infection rate of selected top 10 spreader nodes by SIS model.

| Rank | Proteins | Closeness Centrality | Sum of SIS infection rate of top 10 nodes |
| --- | --- | --- | --- |
| 1 | 1 | 0.085 | 1.94 |
| 2 | 5 | 0.083 |  |
| 3 | 2 | 0.082 |  |
| 4 | 4 | 0.082 |  |
| 5 | 23 | 0.081 |  |
| 6 | 3 | 0.081 |  |
| 7 | 21 | 0.081 |  |
| 8 | 22 | 0.081 |  |
| 9 | 7 | 0.08 |  |
| 10 | 15 | 0.08 |  |
| 11 | 16 | 0.08 | -- |
| 12 | 19 | 0.079 |  |
| 13 | 14 | 0.079801 |  |
| 14 | 9 | 0.079602 |  |
| 15 | 20 | 0.079602 |  |
| 16 | 6 | 0.079602 |  |
| 17 | 8 | 0.079404 |  |
| 18 | 17 | 0.078818 |  |
| 19 | 10 | 0.078624 |  |
| 20 | 11 | 0.078049 |  |
| 21 | 12 | 0.078049 |  |
| 22 | 13 | 0.078049 |  |
| 23 | 18 | 0.07767 |  |
| 24 | 24 | 0.041558 |  |
| 25 | 32 | 0.041237 |  |
| 26 | 28 | 0.041237 |  |
| 27 | 30 | 0.041237 |  |
| 28 | 25 | 0.041237 |  |
| 29 | 27 | 0.041237 |  |
| 30 | 31 | 0.041184 |  |
| 31 | 33 | 0.041184 |  |
| 32 | 29 | 0.040921 |  |
| 33 | 26 | 0.040921 |  |

**Table 3.** Computation of BC of synthetic figure 1 along with computation of infection rate of selected top 10 spreader nodes by SIS model.

| Rank | Proteins | Betweenness Centrality | Sum of SIS infection rate of top 10 nodes |
| --- | --- | --- | --- |
| 1 | 1 | 269.1 | 2.2 |
| 2 | 2 | 117.93 |  |
| 3 | 4 | 117.1 |  |
| 4 | 3 | 114 |  |
| 5 | 5 | 108 |  |
| 6 | 24 | 57 |  |
| 7 | 23 | 56.4 |  |
| 8 | 19 | 45.56 |  |
| 9 | 17 | 39.1 |  |
| 10 | 7 | 36.9 |  |
| 11 | 6 | 32.9 | -- |
| 12 | 18 | 32 |  |
| 13 | 21 | 29.36 |  |
| 14 | 22 | 20.53 |  |
| 15 | 16 | 12.1 |  |
| 16 | 15 | 12.1 |  |
| 17 | 14 | 12.1 |  |
| 18 | 28 | 7 |  |
| 19 | 30 | 7 |  |
| 20 | 25 | 7 |  |
| 21 | 27 | 7 |  |
| 22 | 20 | 6.63 |  |
| 23 | 9 | 4.16 |  |
| 24 | 32 | 1 |  |
| 25 | 29 | 1 |  |
| 26 | 26 | 1 |  |
| 27 | 8 | 0 |  |
| 28 | 11 | 0 |  |
| 29 | 12 | 0 |  |
| 30 | 13 | 0 |  |
| 31 | 10 | 0 |  |
| 32 | 31 | 0 |  |
| 33 | 33 | 0 |  |

**Table 4.** Computation of LAC of synthetic figure 1 along with computation of infection rate of selected top 10 spreader nodes by SIS model.

| Rank | Proteins | Local Average Centrality | Sum of SIS infection rate of top 10 nodes |
| --- | --- | --- | --- |
| 1 | 21 | 2.4 | 2.19 |
| 2 | 9 | 2 |  |
| 3 | 22 | 2 |  |
| 4 | 8 | 2 |  |
| 5 | 11 | 2 |  |
| 6 | 12 | 2 |  |
| 7 | 13 | 2 |  |
| 8 | 2 | 1.6 |  |
| 9 | 23 | 1.6 |  |
| 10 | 7 | 1.5 |  |
| 11 | 3 | 1.5 | -- |
| 12 | 5 | 1.5 |  |
| 13 | 16 | 1.33 |  |
| 14 | 15 | 1.33 |  |
| 15 | 20 | 1.33 |  |
| 16 | 32 | 1.33 |  |
| 17 | 19 | 1 |  |
| 18 | 10 | 1 |  |
| 19 | 31 | 1 |  |
| 20 | 33 | 1 |  |
| 21 | 24 | 0.57 |  |
| 22 | 4 | 0.5 |  |
| 23 | 17 | 0.5 |  |
| 24 | 1 | 0 |  |
| 25 | 14 | 0 |  |
| 26 | 18 | 0 |  |
| 27 | 6 | 0 |  |
| 28 | 28 | 0 |  |
| 29 | 29 | 0 |  |
| 30 | 30 | 0 |  |
| 31 | 25 | 0 |  |
| 32 | 26 | 0 |  |
| 33 | 27 | 0 |  |

**Table 5.** Computation of DC of synthetic figure 1 along with computation of infection rate of selected top 10 spreader nodes by SIS model.

| Rank | Proteins | Degree Centrality | Sum of SIS infection rate of top 10 nodes |
| --- | --- | --- | --- |
| 1 | 24 | 7 | 2.3 |
| 2 | 2 | 5 |  |
| 3 | 23 | 5 |  |
| 4 | 21 | 5 |  |
| 5 | 1 | 4 |  |
| 6 | 7 | 4 |  |
| 7 | 9 | 4 |  |
| 8 | 3 | 4 |  |
| 9 | 4 | 4 |  |
| 10 | 17 | 4 |  |
| 11 | 5 | 4 | -- |
| 12 | 22 | 4 |  |
| 13 | 19 | 4 |  |
| 14 | 8 | 3 |  |
| 15 | 11 | 3 |  |
| 16 | 12 | 3 |  |
| 17 | 13 | 3 |  |
| 18 | 16 | 3 |  |
| 19 | 15 | 3 |  |
| 20 | 20 | 3 |  |
| 21 | 32 | 3 |  |
| 22 | 10 | 2 |  |
| 23 | 14 | 2 |  |
| 24 | 18 | 2 |  |
| 25 | 6 | 2 |  |
| 26 | 28 | 2 |  |
| 27 | 29 | 2 |  |
| 28 | 30 | 2 |  |
| 29 | 25 | 2 |  |
| 30 | 26 | 2 |  |
| 31 | 27 | 2 |  |
| 32 | 31 | 2 |  |
| 33 | 33 | 2 |  |

**Table 6.** Ranking of spreader edges for network 1 and network 2 in Figure 2

| Spreader Edges |  |  |  |  |  |
| --- | --- | --- | --- | --- | --- |
| Rank | Spreader nodes in network 1 | Spreader nodes in network 2 | Spreadability Index of spreader nodes in network 1 | Spreadability Index of spreader nodes in network 2 | Ranking of spreader edges |
| 1 | F | 1 | 5.5 | 76.15 | 40.825 |
| 2 | F | 24 | 5.5 | 22.70 | 14.104 |
| 3 | D | 4 | 5.5 | 15.11 | 10.308 |
| 4 | E | 5 | 4.7 | 13.48 | 9.0919 |
| 5 | E | 19 | 4.7 | 11.66 | 8.1833 |

**Table 7.** Computation of spreadability index of SARS-CoV PPIN along with computation of infection rate of selected top 6 spreader nodes by SIS model.

| Rank | Proteins | $E_{out}^{S_i}$ | $E_{in}^{S_i}$ | Edge Ratio | Neighborhood Density | Node Weight | Spreadability Index | SIS infection rate of top 6 nodes | Sum of SIS infection rate of top 6 nodes |
| --- | --- | --- | --- | --- | --- | --- | --- | --- | --- |
| 1 | M | 7 | 3 | 2.0 | 3.845 | 3.4 | 11.090 | 1 | 2.935 |
| 2 | S | 6 | 3 | 1.75 | 4.047 | 3.2 | 10.283 | 0.2 |  |
| 3 | ORF8AB | 7 | 3 | 2.0 | 1.785 | 4.0 | 7.5714 | 1 |  |
| 4 | ORF8B | 5 | 5 | 1.0 | 3.464 | 3.8 | 7.2642 | 0.2 |  |
| 5 | E | 7 | 3 | 2.0 | 1.428 | 4.0 | 6.8571 | 0.25 |  |
| 6 | ORF3A | 2 | 8 | 0.333 | 9.249 | 3.428 | 6.5119 | 0.285 |  |
| 7 | ORF7A | 2 | 8 | 0.333 | 9.25 | 3.428 | 6.5119 | -- | -- |
| 8 | ORF8A | 3 | 0 | 4.0 | 0.0 | 2.0 | 2 |  |  |
| 9 | N | 3 | 0 | 4.0 | 0.0 | 2.0 | 2 |  |  |

**Table 8.** Computation of degree centrality of SARS-CoV PPIN along with computation of infection rate of selected top 6 spreader nodes by SIS model.

| Rank | Proteins | Degree Centrality [19] | SIS infection rate of top 6 nodes | Sum of SIS infection rate of top 6 nodes |
| --- | --- | --- | --- | --- |
| 1 | ORF7A | 6 | 0.285 | 1.82 |
| 2 | ORF3A | 6 | 0.285 |  |
| 3 | ORF8B | 4 | 0.2 |  |
| 4 | M | 4 | 0.6 |  |
| 5 | S | 4 | 0.2 |  |
| 6 | E | 3 | 0.25 |  |
| 7 | ORF8AB | 3 | -- | -- |
| 8 | N | 1 |  |  |
| 9 | ORF8A | 1 |  |  |

**Table 9.** Computation of closeness centrality of SARS-CoV PPIN along with computation of infection rate of selected top 6 spreader nodes by SIS model.

| Rank | Proteins | Closeness Centrality [24] | SIS infection rate of top 6 nodes | Sum of SIS infection rate of top 6 nodes |
| --- | --- | --- | --- | --- |
| 1 | ORF7A | 0.239 | 0.285 | 1.82 |
| 2 | ORF3A | 0.239 | 0.285 |  |
| 3 | ORF8B | 0.224 | 0.2 |  |
| 4 | M | 0.224 | 0.6 |  |
| 5 | S | 0.224 | 0.2 |  |
| 6 | E | 0.215 | 0.25 |  |
| 7 | ORF8AB | 0.22 |  |  |
| 8 | N | 0.196 | -- | -- |
| 9 | ORF8A | 0.196 |  |  |

**Table 10.** Computation of local average centrality of SARS-CoV PPIN along with computation of infection rate of selected top 6 spreader nodes by SIS model.

| Rank | Proteins | Local Average Centrality [21] | SIS infection rate of top 6 nodes | Sum of SIS infection rate of top 6 nodes |
| --- | --- | --- | --- | --- |
| 1 | ORF7A | 2.666 | 0.285 | 2.22 |
| 2 | ORF3A | 2.666 | 0.285 |  |
| 3 | ORF8B | 2.5 | 0.2 |  |
| 4 | E | 2 | 0.25 |  |
| 5 | ORF8AB | 2 | 1 |  |
| 6 | S | 1.5 | 0.2 |  |
| 7 | M | 1.5 | -- | -- |
| 8 | N | 0 |  |  |
| 9 | ORF8A | 0 |  |  |

**Table 11.**
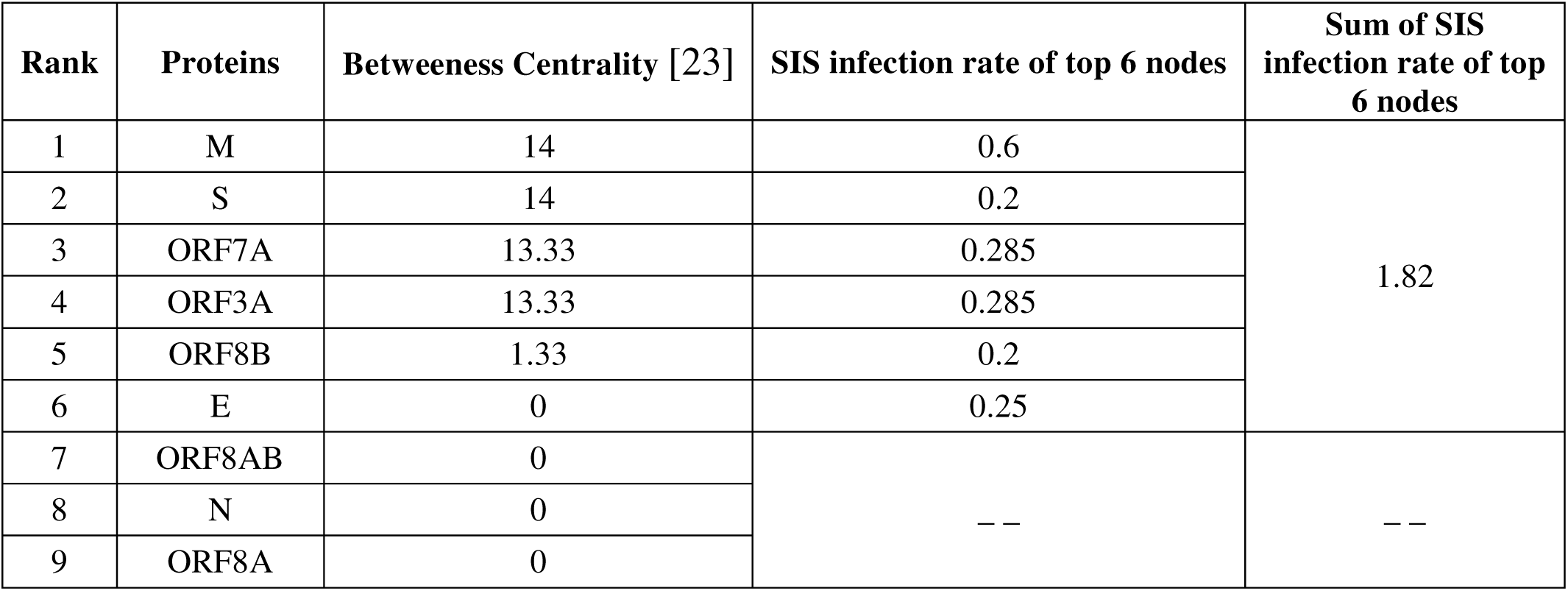
Computation of betweeness centrality of SARS-CoV PPIN along with computation of infection rate of selected top 6 spreader nodes by SIS model.

**Table 12.** Network statistics of spreaders at three levels of thresholds

| Threshold | SARS-CoV spreaders | SARS-CoV-s level 1 human spreaders | SARS-CoV-s level 2 human spreaders |
| --- | --- | --- | --- |
| High | 6 | 24 | 9 |
| Medium | 6 | 24 | 22 |
| Low | 6 | 24 | 111 |

**Table 13.**
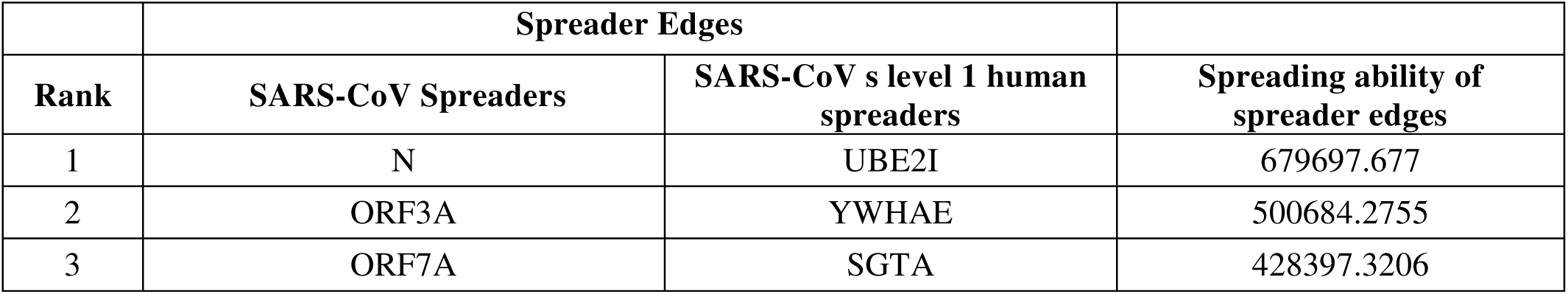
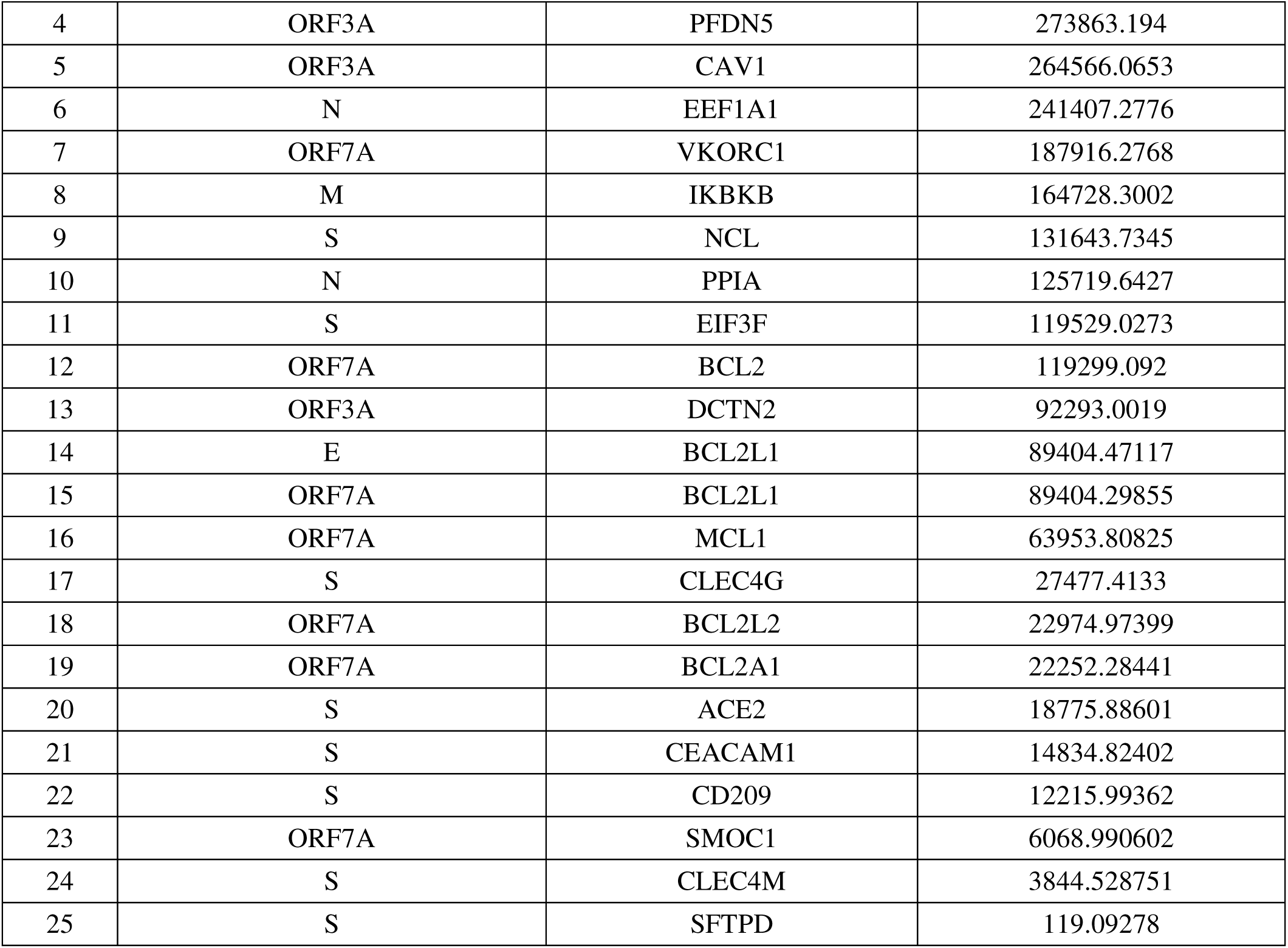
Ranked spreader edges between SARS-CoV spreaders and its level 1 human spreaders

The entire network view of SARS-CoV and human PPIN has been generated online under three circumstances:

1. All the nodes and edges are considered as spreader nodes and edges respectively and are ranked accordingly. https://yu2qkp7gwoinjwsebyw0xw-on.drv.tw/www.graph_all.html/graph_all.html
2. Selected Spreader nodes and edges are highlighted for high threshold. https://yu2qkp7gwoinjwsebyw0xw-on.drv.tw/www.high_threshold.com/graph_high_threshold.html
3. Selected Spreader nodes and edges are highlighted for medium threshold. https://yu2qkp7gwoinjwsebyw0xw-on.drv.tw/www.medium_threshold.com/graph_medium_threshold.html
4. Selected Spreader nodes and edges are highlighted for low threshold. https://yu2qkp7gwoinjwsebyw0xw-on.drv.tw/www.low_threshold.com/graph_low_threshold.html

In the above generated network views, the blue, yellow and green color represent SARS-CoV spreaders, its level 1 human spreaders and its level2 human spreaders respectively. The remaining nodes are in indigo.

## 4. Conclusion

Spreadability index is thus proved to be effective in the detection of spreader nodes and edges in SARS-CoV-human PPIN along with the cross validation by SIS model. Spreader nodes are the most critical nodes in the network which actually transmit the infection to its successors. Simultaneously it is also true if the spreader nodes are not connected with spreader edges infection transmission would not have been possible.

In a nutshell, it can be said that the proposed work exploits the possibility of understanding the entire disease propagation/transmission from SARS-CoV network to human network. It should be borne in mind that SARS-CoV2 is ∼89% genetically similar to its predecessor SARS-CoV [34], [35]. It strongly reveals the fact that the human proteins chosen as spreaders of SARS-CoV might be the potential targets of SARS-CoV 2 also. If our future research work reveals this assumption, then it will definitely explore a new direction in the identification of essential drugs/vaccine for SARS-CoV2.

## Acknowledgement

Authors are thankful to the CMATER research laboratory of the Computer Science Department, Jadavpur University, India, for providing infrastructure facilities during progress of the work. This project is partially supported by the CMATER research laboratory of the Computer Science and Engineering Department, Jadavpur University, India, and DBT project (No.BT/PR16356/BID/7/596/2016), Ministry of Science and Technology, Government of India.

## Appendix

**A-Table 1.** Ranked spreader edges between SARS-CoV s level 1 and level 2 human spreaders at high threshold

|  | Spreader Edges |  |  |
| --- | --- | --- | --- |
| Rank | SARS-CoV s level 1 and level 2 human spreaders |  | Spreading ability of spreader edges |
| 1 | APP | UBE2I | 9297903 |
| 2 | APP | CAV1 | 8882769 |
| 3 | APP | PPIA | 8743925 |
| 4 | APP | EIF3F | 8737730 |
| 5 | BCL2A1 | APP | 8640455 |
| 6 | ELAVL1 | EEF1A1 | 4284371 |
| 7 | NCL | ELAVL1 | 4174604 |
| 8 | PPIA | ELAVL1 | 4168684 |
| 9 | BCL2L2 | ELAVL1 | 4065937 |
| 10 | NXF1 | UBE2I | 3653905 |
| 11 | NXF1 | SGTA | 3402603 |
| 12 | NTRK1 | YWHAE | 3275095 |
| 13 | NXF1 | CAV1 | 3238771 |
| 14 | NXF1 | EEF1A1 | 3215615 |
| 15 | XPO1 | UBE2I | 3169805 |
| 16 | DCTN2 | NXF1 | 3066498 |
| 17 | NTRK1 | CAV1 | 3038977 |
| 18 | NXF1 | MCL1 | 3038159 |
| 19 | NTRK1 | EEF1A1 | 3015820 |
| 20 | XPO1 | YWHAE | 2990790 |
| 21 | NCL | NTRK1 | 2906052 |
| 22 | NTRK1 | PPIA | 2900133 |
| 23 | NTRK1 | EIF3F | 2893938 |
| 24 | DCTN2 | NTRK1 | 2866704 |
| 25 | GRB2 | UBE2I | 2349446 |
| 26 | GRB2 | YWHAE | 2170431 |
| 27 | IKBKB | MEOX2 | 2061694 |
| 28 | UBC | SGTA | 2048943 |
| 29 | YWHAE | EGFR | 2023580 |
| 30 | GRB2 | PFDN5 | 1943610 |
| 31 | GRB2 | EEF1A1 | 1911156 |

**A-Table 2.** Ranked spreader edges between SARS-CoV s level 1 and level 2 human proteins at medium threshold

|  | Spreader Edges |  |  |
| --- | --- | --- | --- |
| Rank | SARS-CoV s level 1 and level 2 human spreaders |  | Spreading ability of spreader edges |
| 1 | APP | UBE2I | 9297902.664 |
| 2 | APP | CAV1 | 8882768.796 |
| 3 | APP | PPIA | 8743924.63 |
| 4 | APP | EIF3F | 8737729.873 |
| 5 | BCL2A1 | APP | 8640455.016 |
| 6 | ELAVL1 | EEF1A1 | 4284371.281 |
| 7 | NCL | ELAVL1 | 4174603.596 |
| 8 | PPIA | ELAVL1 | 4168683.646 |
| 9 | BCL2L2 | ELAVL1 | 4065936.721 |
| 10 | NXF1 | UBE2I | 3653905.316 |
| 11 | NXF1 | SGTA | 3402602.703 |
| 12 | NTRK1 | YWHAE | 3275094.903 |
| 13 | NXF1 | CAV1 | 3238771.448 |
| 14 | NXF1 | EEF1A1 | 3215614.916 |
| 15 | XPO1 | UBE2I | 3169805.311 |
| 16 | DCTN2 | NXF1 | 3066498.385 |
| 17 | NTRK1 | CAV1 | 3038976.693 |
| 18 | NXF1 | MCL1 | 3038159.191 |
| 19 | NTRK1 | EEF1A1 | 3015820.161 |
| 20 | XPO1 | YWHAE | 2990789.654 |
| 21 | NCL | NTRK1 | 2906052.476 |
| 22 | NTRK1 | PPIA | 2900132.526 |
| 23 | NTRK1 | EIF3F | 2893937.769 |
| 24 | DCTN2 | NTRK1 | 2866703.629 |
| 25 | GRB2 | UBE2I | 2349446.25 |
| 26 | GRB2 | YWHAE | 2170430.592 |
| 27 | IKBKB | MEOX2 | 2061694.24 |
| 28 | UBC | SGTA | 2048942.842 |
| 29 | YWHAE | EGFR | 2023579.783 |
| 30 | GRB2 | PFDN5 | 1943609.51 |
| 31 | GRB2 | EEF1A1 | 1911155.85 |
| 32 | UBC | EEF1A1 | 1861955.055 |
| 33 | TP53 | UBE2I | 1829375.898 |
| 34 | BAG3 | UBE2I | 1809920.855 |
| 35 | NCL | GRB2 | 1801388.165 |
| 36 | GRB2 | PPIA | 1795468.215 |
| 37 | CAV1 | EGFR | 1787461.572 |
| 38 | GOLGA2 | UBE2I | 1779690.372 |
| 39 | EEF1A1 | EGFR | 1764305.041 |
| 40 | DCTN2 | GRB2 | 1762039.318 |
| 41 | NCL | UBC | 1752187.37 |
| 42 | EIF3F | UBC | 1740072.663 |
| 43 | YWHAE | SHMT2 | 1738748.567 |
| 44 | DCTN2 | UBC | 1712838.523 |
| 45 | BCL2A1 | GRB2 | 1691998.601 |
| 46 | MCL1 | UBC | 1684499.329 |
| 47 | TP53 | YWHAE | 1650360.24 |
| 48 | EIF3F | EGFR | 1642422.649 |
| 49 | PROP1 | UBE2I | 1641256.177 |
| 50 | BAG3 | YWHAE | 1630905.197 |
| 51 | CDK2 | UBE2I | 1622979.319 |
| 52 | DCTN2 | EGFR | 1615188.509 |
| 53 | CUL3 | YWHAE | 1614438.165 |
| 54 | TRIM27 | UBE2I | 1579133.993 |
| 55 | CEACAM1 | EGFR | 1537728.445 |
| 56 | HSP90AA1 | YWHAE | 1503676.376 |
| 57 | TRAF6 | UBE2I | 1451472.666 |
| 58 | CDK2 | YWHAE | 1443963.661 |
| 59 | MCM2 | YWHAE | 1440253.354 |
| 60 | BRCA1 | UBE2I | 1439839.399 |
| 61 | HSP90AA1 | SGTA | 1431389.421 |
| 62 | ESR1 | UBE2I | 1424801.897 |

**A-Table 3.**
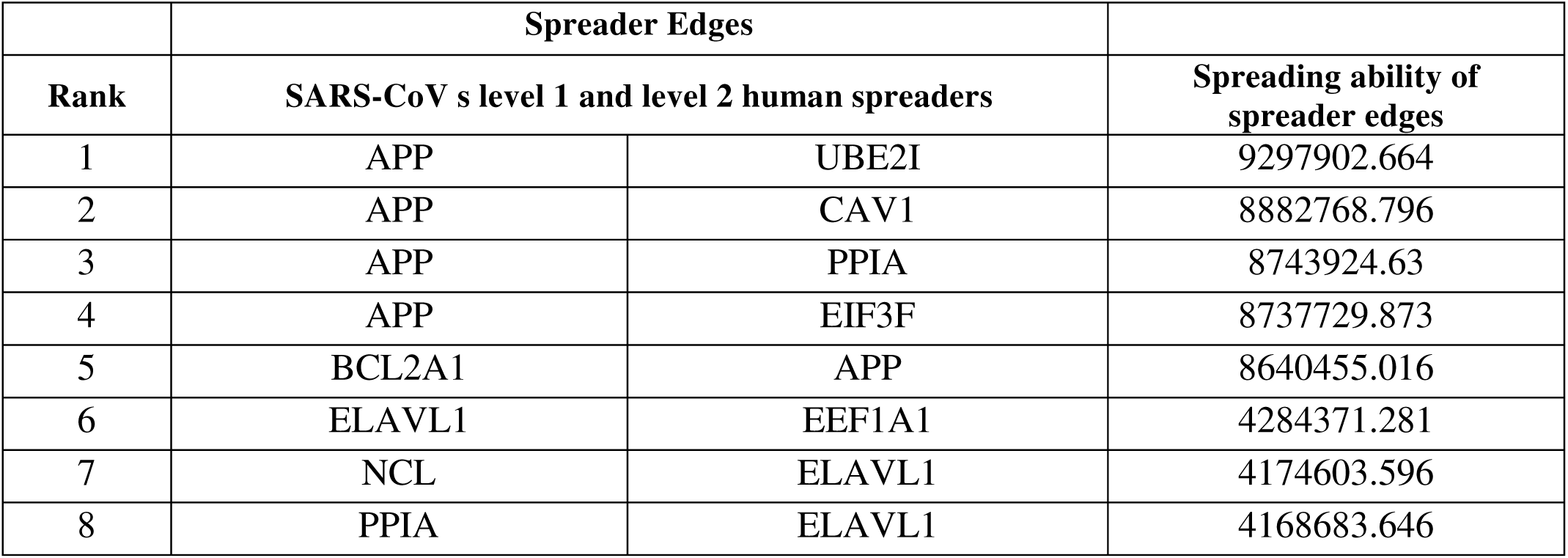
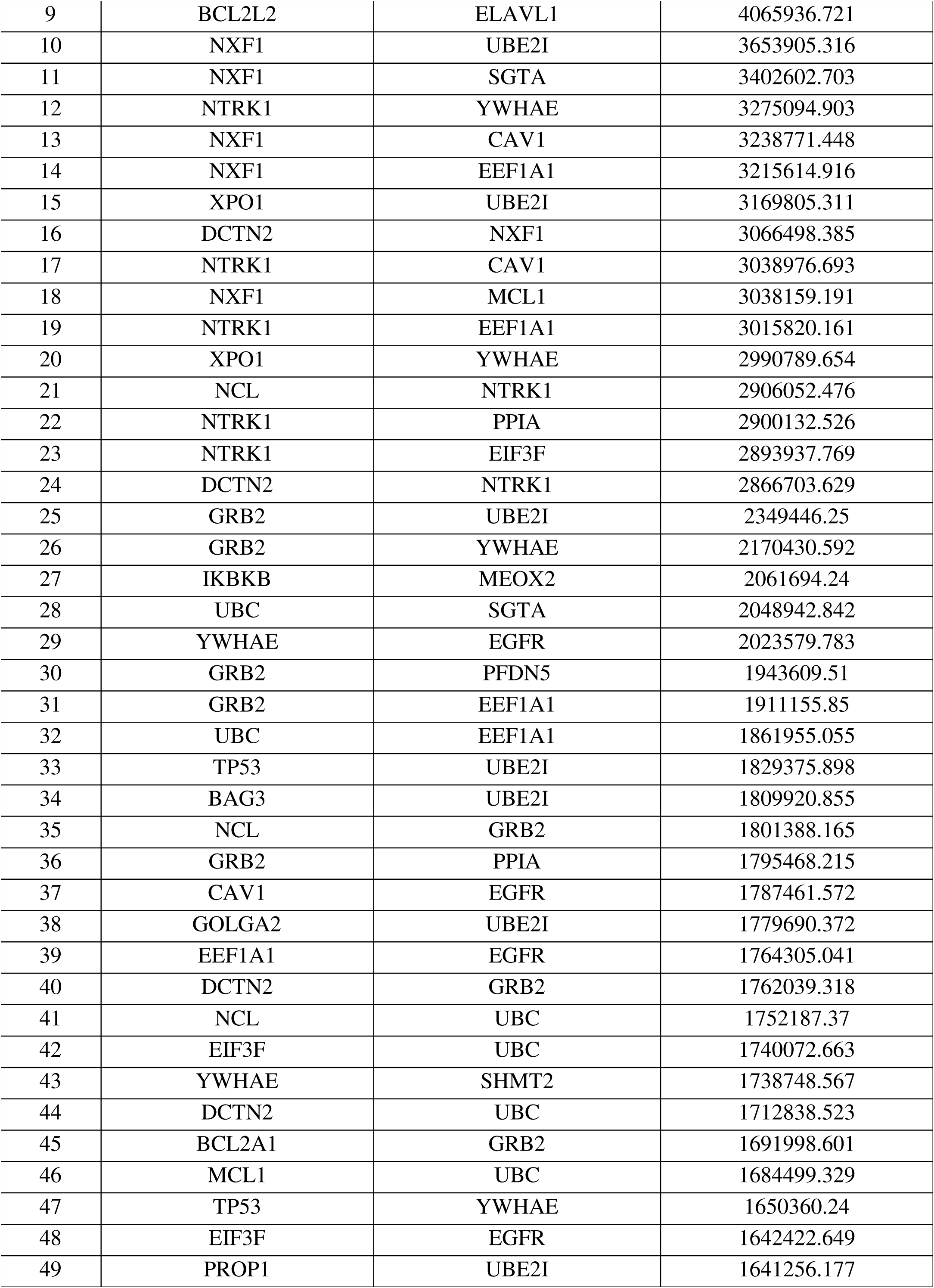
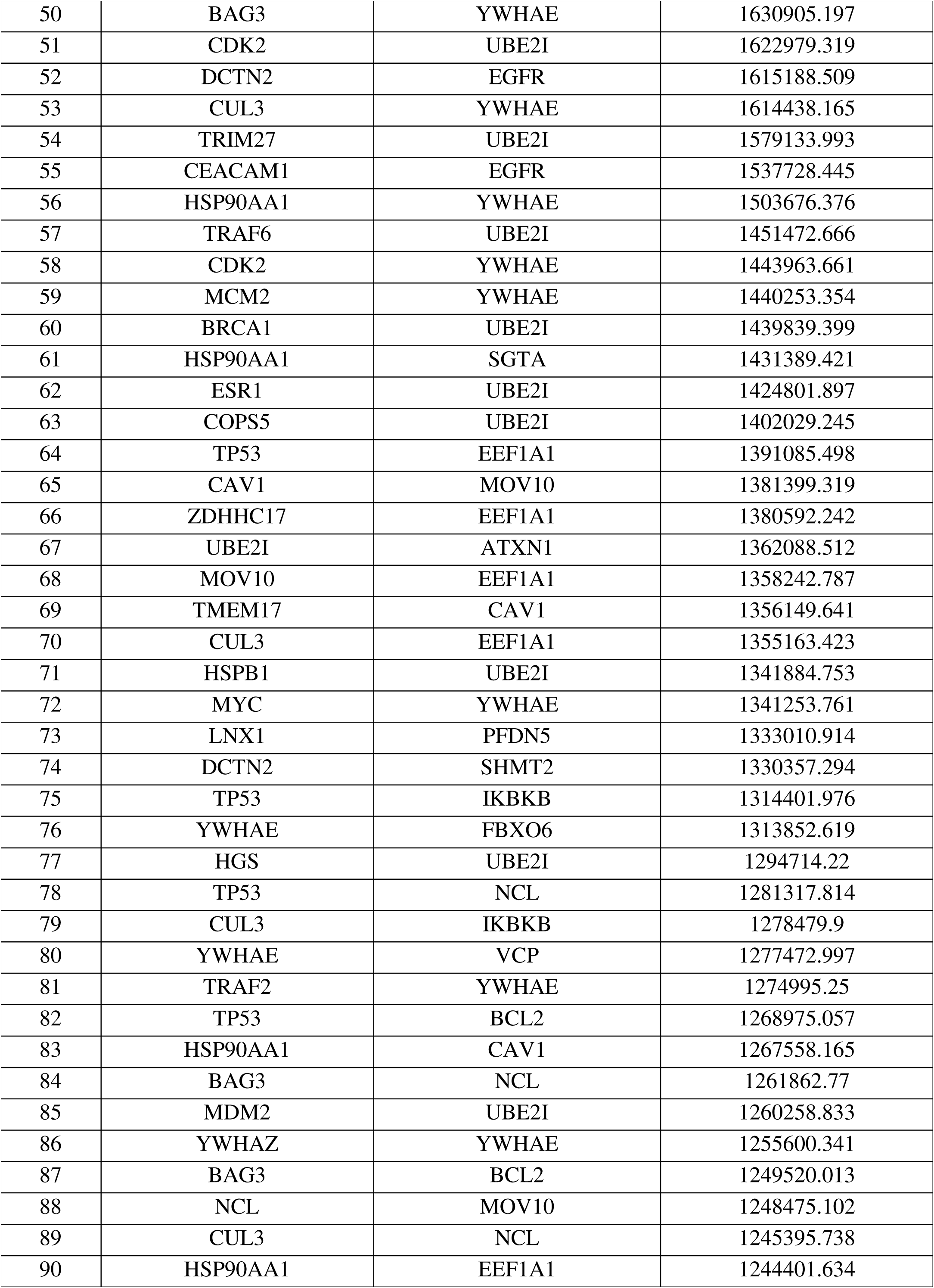
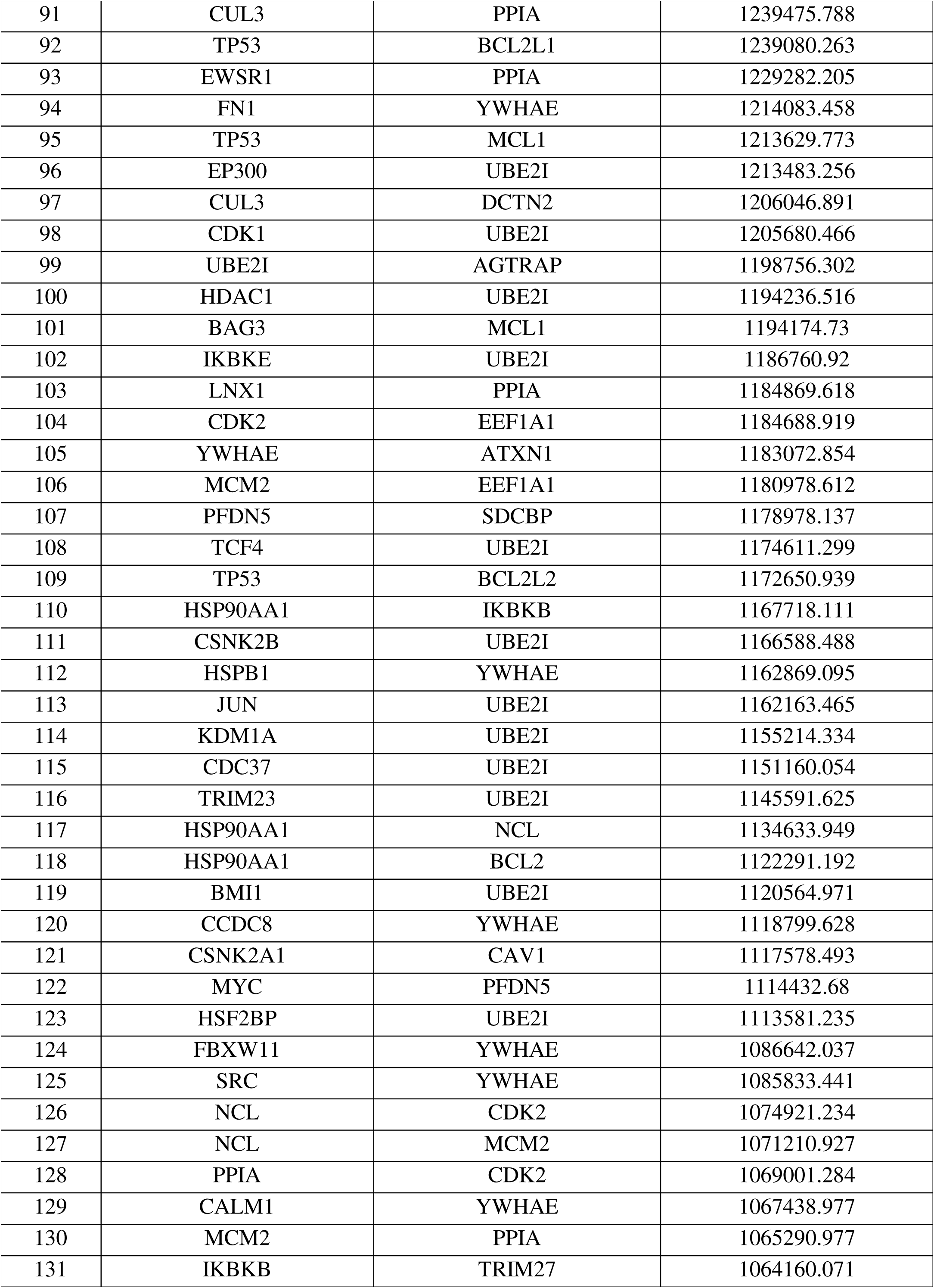
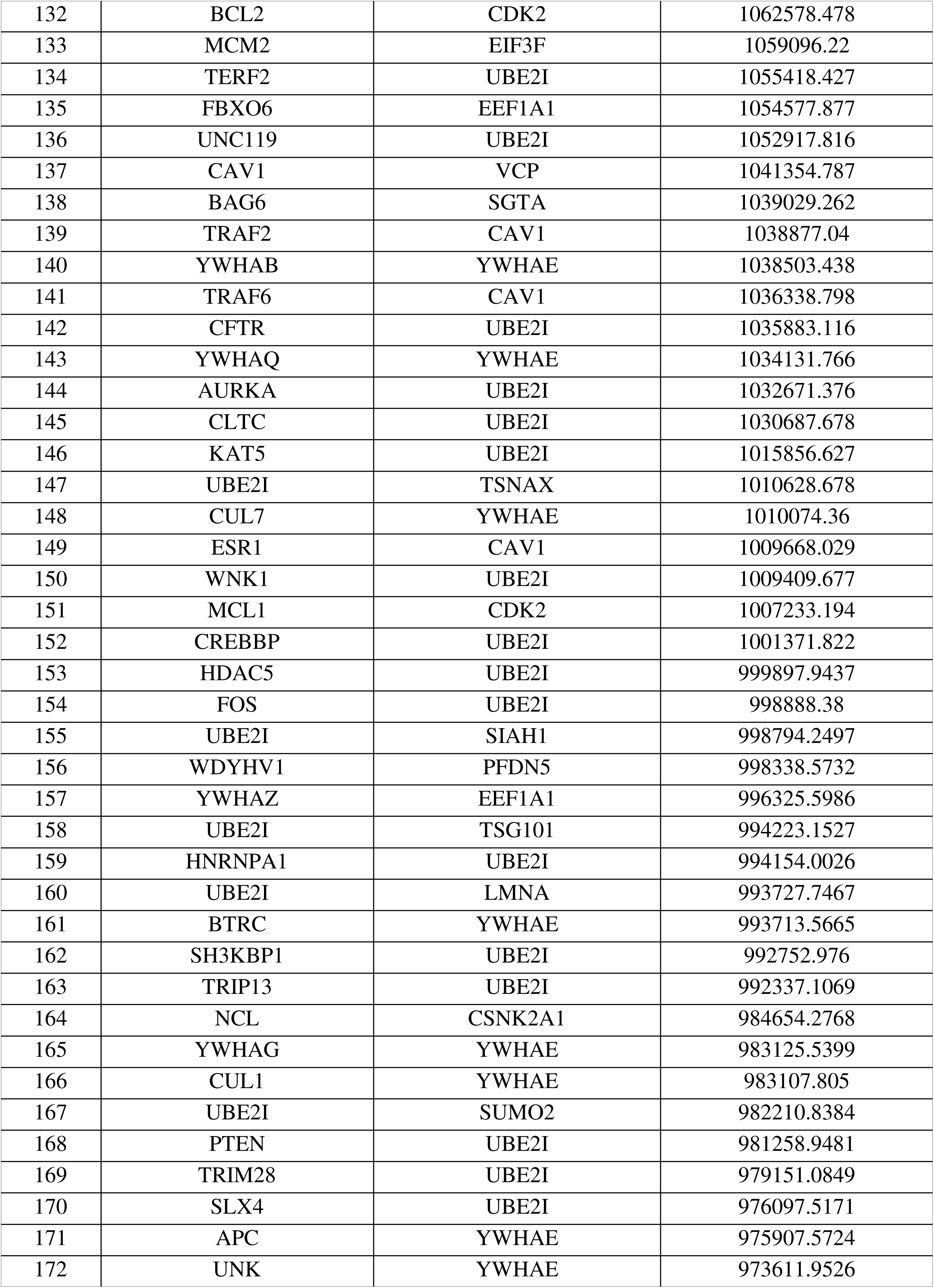
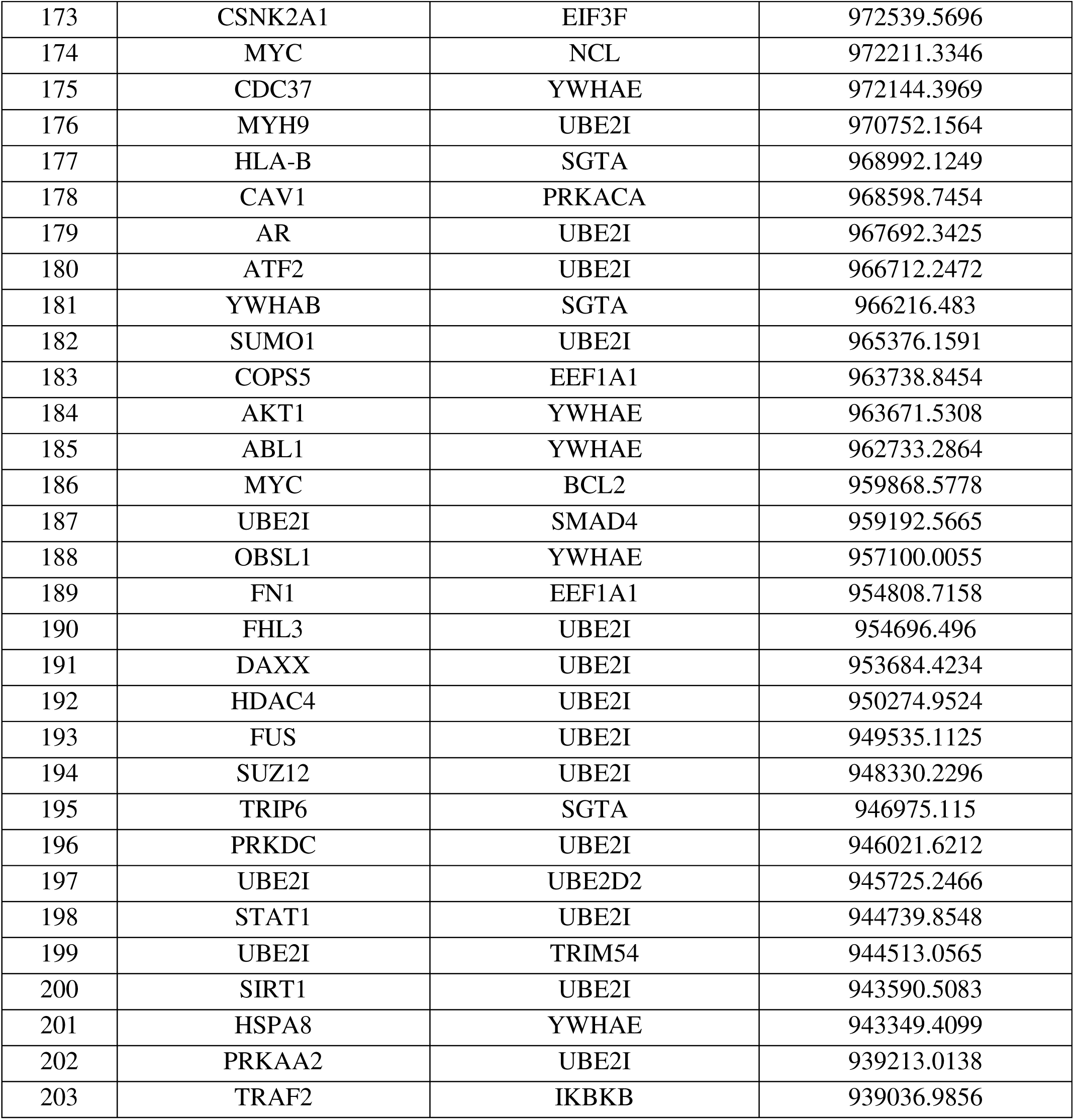
Ranked spreader edges between SARS-CoV s level 1 and level 2 human spreaders at low threshold

